# Rap1 binding and a lipid-dependent helix in talin F1 domain promote integrin activation in tandem

**DOI:** 10.1101/504894

**Authors:** Alexandre R. Gingras, Frederic Lagarrigue, Monica N. Cuevas, Andrew J. Valadez, Marcus Zorovich, Wilma McLaughlin, Miguel Alejandro Lopez-Ramirez, Nicolas Seban, Klaus Ley, William B. Kiosses, Mark H. Ginsberg

## Abstract

Rap1 GTPases bind effectors, such as RIAM, to enable talin1 to induce integrin activation. In addition, Rap1 binds directly to the talin1 F0 domain (F0); however, this interaction makes a limited contribution to integrin activation in CHO cells or platelets. Here, we show that talin1 F1 domain contains a previously undetected Rap1 binding site of similar affinity to that in F0. A structure-guided point mutant (R118E) in F1, which blocks Rap1 binding, abolishes the capacity of Rap1 to potentiate talin1-induced integrin activation. The capacity of F1 to mediate Rap1-dependent integrin activation depends on a unique loop in F1 that transforms into an helix upon binding to membrane lipids. Basic membrane-facing residues of this helix are critical as charge reversal mutations led to dramatic suppression of talin1-dependent activation. Thus, a novel Rap1 binding site and a lipid-dependent helix in talin1 F1 work in tandem to enable a direct Rap1-talin1 interaction to cause integrin activation.

**Summary:** This work reveals that Rap1 GTPases bind directly to talin1 F1 domain and by cooperating with a unique lipid-dependent amphipathic helix in the F1 domain effects lin1-mediated integrin activation.

## Introduction

Integrin receptors are critical mediators of cellular adhesion, migration and assembly of the extracellular matrix, thereby playing an indispensable role in development and in many pathological processes (Hynes, 2002). Regulation of the affinity of integrins for their ligands is central to their function. In particular, integrins in blood cells are usually expressed in a low-affinity form until intracellular signals initiated by agonists acting via distinct excitatory receptors induces a high-affinity state, a process operationally defined as integrin activation. Binding of talin1 to the cytoplasmic tail of integrin β1 (Shattil et al., 2010; Tadokoro et al., 2003), β2 (Simonson et al., 2006), β3 (Haling et al., 2011; Nieswandt et al., 2007; Petrich et al., 2007a; Petrich et al., 2007b), and β7 (Sun et al., 2018) is a critical final common step in integrin activation.

Talin1 is a large (270 kDa) multi-domain protein that links integrins to the actin cytoskeleton via its N-terminal head domain that binds to the β-integrin cytoplasmic tail and its C-terminal flexible rod domain that binds F-actin (Critchley and Gingras, 2008). The talin1 head domain (THD) comprises an atypical FERM domain because it contains an additional F0 domain to the characteristic FERM F1, F2 and F3 domains, and because it can adopt a linear arrangement rather than the cloverleaf structure typically observed in other FERM domains (Calderwood et al., 2013). Talin1 is auto-inhibited in the cytosol due to the interaction of the talin1 head domain (THD) with the rod domain, which prevents the interaction of THD with the integrin β cytoplasmic tail (Song et al., 2012). Our understanding of the signaling events that regulate talin1 recruitment to the plasma membrane and its association with integrin is incomplete.

Rap1 GTPases are perhaps the most completely studied relays of signals from cell surface receptors to integrin activation. Combined deficiency of both Rap1A and Rap1B isoforms in the megakaryocyte lineage evidenced an essential role for Rap1 signaling in platelet integrin activation and function (Stefanini et al., 2018). Although the Rap1 effector RIAM plays a major role in recruiting talin1 in leukocytes (Klapproth et al., 2015; Lagarrigue et al., 2017; Su et al., 2015), its role in talin1-dependent activation of platelet integrins has been unequivocally ruled out (Klapproth et al., 2015; Stritt et al., 2015; Su et al., 2015).

Studies using recombinant αIIbβ3-expressing A5 CHO cells revealed that Rap1 activity regulates talin1 head domain (THD)-induced integrin αIIbβ3 activation (Lagarrigue et al., 2018), whilst activation by the F2F3 subdomain is Rap1-independent (Han et al., 2006). This result suggests that a direct interaction between Rap1 and the F0F1 subdomain domain facilitates integrin activation. Structural studies showed that Rap1b binds directly to talin1 F0 domain with low affinity (Goult et al., 2010). Furthermore, in *Dictyostelium* Rap1 directly interacts with the RA domain of talinB (Plak et al., 2016) to enable adhesion during *Dictyostelium* morphogenesis. Similarly, a mutation in F0 that inhibits Rap1 binding is embryonic lethal in Drosophila (Camp et al., 2018). Zhu et al. confirmed the direct Rap1-talin1 F0 interaction and showed that membrane-anchored Rap1b in vesicles has enhanced binding to THD, suggesting a mechanism of talin1 recruitment to integrins by Rap1 (Zhu et al., 2017). Quantitative proteomic analyses of murine platelets revealed the high abundance of Rap1b and talin1 (Zeiler et al., 2014). The abundance of the proteins at equal molar ratios and the lack of a known Rap1 effector with such a high abundance in platelets suggest that talin1 F0 domain may act as a direct effector of Rap1 to activate integrins in platelets. However, we recently reported a talin1 point mutation (R35E) in F0 domain that reduces Rap1 affinity by greater than 25 fold does not impair the capacity of THD to activate integrins in A5 cells nor does it abolish effects of Rap1 activity on THD-induced activation. In accord with this result, we and others (Bromberger et al., 2018) found that mice with F0 mutations that disrupt Rap1 binding are viable, fertile, apparently healthy, and exhibit similar or slightly reduced extent and kinetics of αIIbβ3 activation (Lagarrigue et al., 2018). These mice did manifest a mild defect in platelet aggregation and in hemostasis. Thus, the low affinity Rap1-talin1 F0 interaction does not make a major contribution to integrin αIIbβ3 activation in mice and cannot account for the profound effects on activation in RIAM-deficient platelets of: i) loss of Rap1 activity (Stefanini et al., 2018); ii) deletion of talin1 (Nieswandt et al., 2007; Petrich et al., 2007b); iii) mutations of integrin β3 that block talin1 binding (Petrich et al., 2007a); and iv) mutations of talin1 that prevents binding to integrin β3 (Haling et al., 2011).

Here, we show that talin1 F1 domain contains a previously undiscovered Rap1 binding site of similar affinity to that in F0. A structure-guided mutant in F1, which blocks Rap1 binding, abolishes the capacity of Rap1 to potentiate talin1-induced integrin activation. Talin1 F1’s ability to mediate Rap1-dependent integrin activation requires a unique unstructured loop that transforms into an amphipathic helix upon binding to membrane lipids. Charge-reversal mutations of basic membrane-facing residues of this helix profoundly suppressed talin1-dependent activation. Thus, a novel Rap1 binding site and a lipid-dependent amphipathic helix in talin1 F1 work in tandem to enable a direct Rap1-talin1 interaction to promote integrin activation.

## Results and Discussion

### THD contains a second Rap1 binding site

Since integrin activation induced by THD(R35E) was still Rap1-dependent (Lagarrigue et al., 2018), we hypothesized that THD could have a second Rap1-binding site. We used NMR spectroscopy to examine the interaction of ^15^N-labelled Rap1b with unlabeled THD. The 2D-HSQC of Rap1b showed dispersed resonances characteristic of a well-folded protein (Fig. 1 A). Addition of THD caused extensive broadening of Rap1b resonances (Fig. 1 B), due to its interaction with the larger THD. We previously showed that the F0 domain of talin1(R35E) does not detectably bind Rap1b (Lagarrigue et al., 2018); however, THD(R35E) caused broadening of the Rap1b resonances (Fig. 1 C), whereas, wild-type THD (Fig. 1 B) caused a more profound effect. While R35E mutation reduces the affinity dramatically, residual binding might still be able to cause broadening. We therefore tested THD(ΔF0), comprising talin1 F1-F3 domains, and found that it bound Rap1 as assessed by intermediate broadening (Fig. 1 D), consistent with the reduced mass of THD(ΔF0). The residual Rap1 binding capacity of THD(R35E) and THD(ΔF0) combined with the lack of Rap1-dependence of F2F3 mediated integrin activation (Han et al., 2006), raised the possibility that THD has a second Rap1 binding site in the F1 domain.

**Figure 1.**
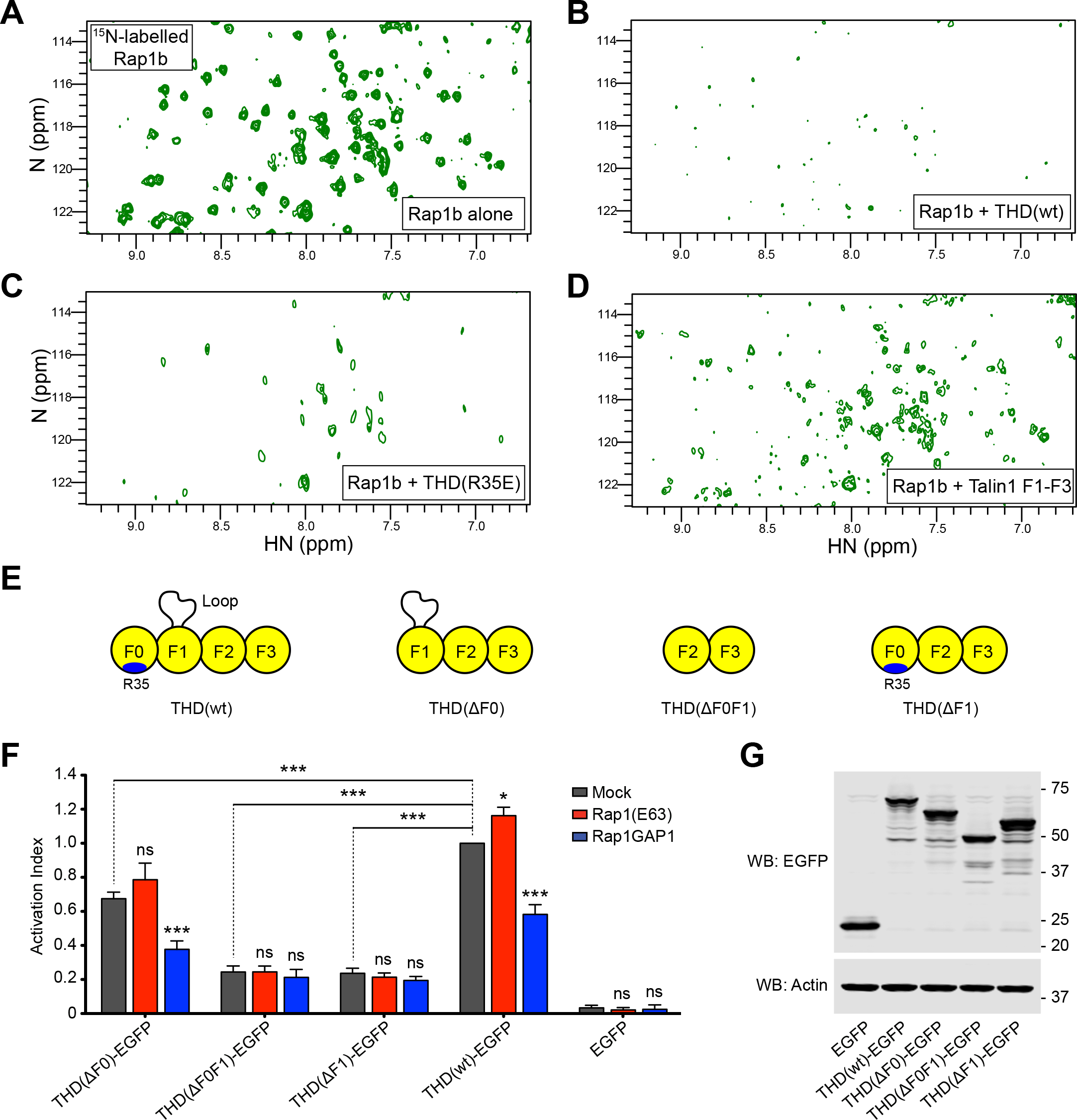
THD contains a second Rap1 binding site. **(A-D)** 2D ^1^H,^15^N-sfHMQC spectra of 100 μM ^15^N-labeled Rap1b. **(A)** Free Rap1b. **(B)** Rap1b with 250 μM THD wild-type. The broadening of Rap1 resonances in the presence of THD indicates binding. **(C)** Rap1b with 250 μM THD(R35E). The broadening indicates the presence of a second Rap1 binding site. **(D)** Rap1b with 250 μM THD(ΔF0). The intermediate broadening indicates the a second Rap1 binding site outside the F0 domain. **(E)** Talin1 head domain (THD) constructs used in panels F and G. **(F)** A5 cells stably expressing αIIbβ3 integrin were transfected with cDNA encoding THD-EGFP in combination with Rap1a(Q63E) or Rap1GAP1. Integrin activation was assayed by binding of PAC1 to EGFP positive cells. Transfection of EGFP alone was used as control. Bar graphs represent Mean ± SEM of 4 independent experiments, and normalized to THD(ΔF0)– EGFP + Mock. Two-way ANOVA with Bonferroni post-test. Each condition was compared to the respective Mock control. ns, not significant; *, p<0.05; ***, pμ0.001. **(G)** Western blot of THD-EGFP expression. Actin was used as a loading control.

To assess the contribution of each subdomain in THD in Rap1-regulated integrin activation, we generated THD deletions by removing either the F0, F1, or F0F1 domains (Fig. 1 E) and tested effects on integrin activation in A5 cells. THD induced a similar extent of integrin activation to addition of manganese (Fig. S1 A) indicating that THD expression provokes substantial integrin αIIbβ3 activation in A5 cells. Expression of all three THD mutants induced activation (Fig. 1 F, grey bars), with a markedly lower activation index for THD(ΔF1) and THD(ΔF0F1), both lacking the F1 domain. Inhibition of endogenous Rap1 activity by co-expression of Rap1GAP1 lead to a decrease in both THD and THD(ΔF0)-mediated activation, but did not affect activation with THD mutants lacking the F1 domain (Fig. 1 F, blue bars). Furthermore, expression of the constitutively activated Rap1b(Q63E) mutant trended to increased THD(ΔF0)-mediated αIIbβ3 activation, but had no evident effect on activation with THD mutants lacking the F1 domain (Fig. 1 F, red bars). Thus, the effect of Rap1 activity on integrin activation by THD(ΔF0) resembles that observed with THD (Lagarrigue et al., 2018). Immunoblotting confirmed similar level of expression for all THD mutants (Fig. 1 G). Because (Bromberger et al., 2018) reported a slight reduction in platelet αIIbβ3 activation by introducing the three mutations, K15A, R30A and R35A, in the F0 domain, we tested the effect of these mutations on THD-dependent αIIbβ3 integrin activation in A5 cells and observed no effect (Fig. S1 B-E). Taken together, these data suggest that talin1 F1 has a second Rap1-binding site important in integrin activation; a surprising conclusion in light of a previous report that the F1 domain does not bind Rap1b (Zhu et al., 2017).

### Talin1 F1 domain binds Rap1

Since both the talin1 F0 and F1 domains have a ubiquitin fold (Goult et al., 2010), we aligned their amino-acid sequence and noticed that the key positively charged residues for Rap1 binding are conserved: K15 and R35 in F0, and R98 and R118 in F1 (Fig. 2 A). Moreover, superimposition of the NMR structures of the talin1 F0 domain with the F1 domain showed that the position of these residues is maintained. We therefore generated a model of the talin1 F1-Rap1 complex by superimposing the talin1 F1 domain with the F0 domain in complex with Rap1b (Zhu et al., 2017). We found that the F1 interface with Rap1b to be very similar to that of F0 with positively charged R98 and R118 showing surface charge complementarity with the negatively charged groups on Rap1 surface (Fig. 2 B). Importantly, using the NMR structure of the talin1 F1 domain alone (Goult et al., 2010), we found the inserted F1 loop to be positioned far away from the binding interface where it would not sterically hinder Rap1b binding.

**Figure 2.**
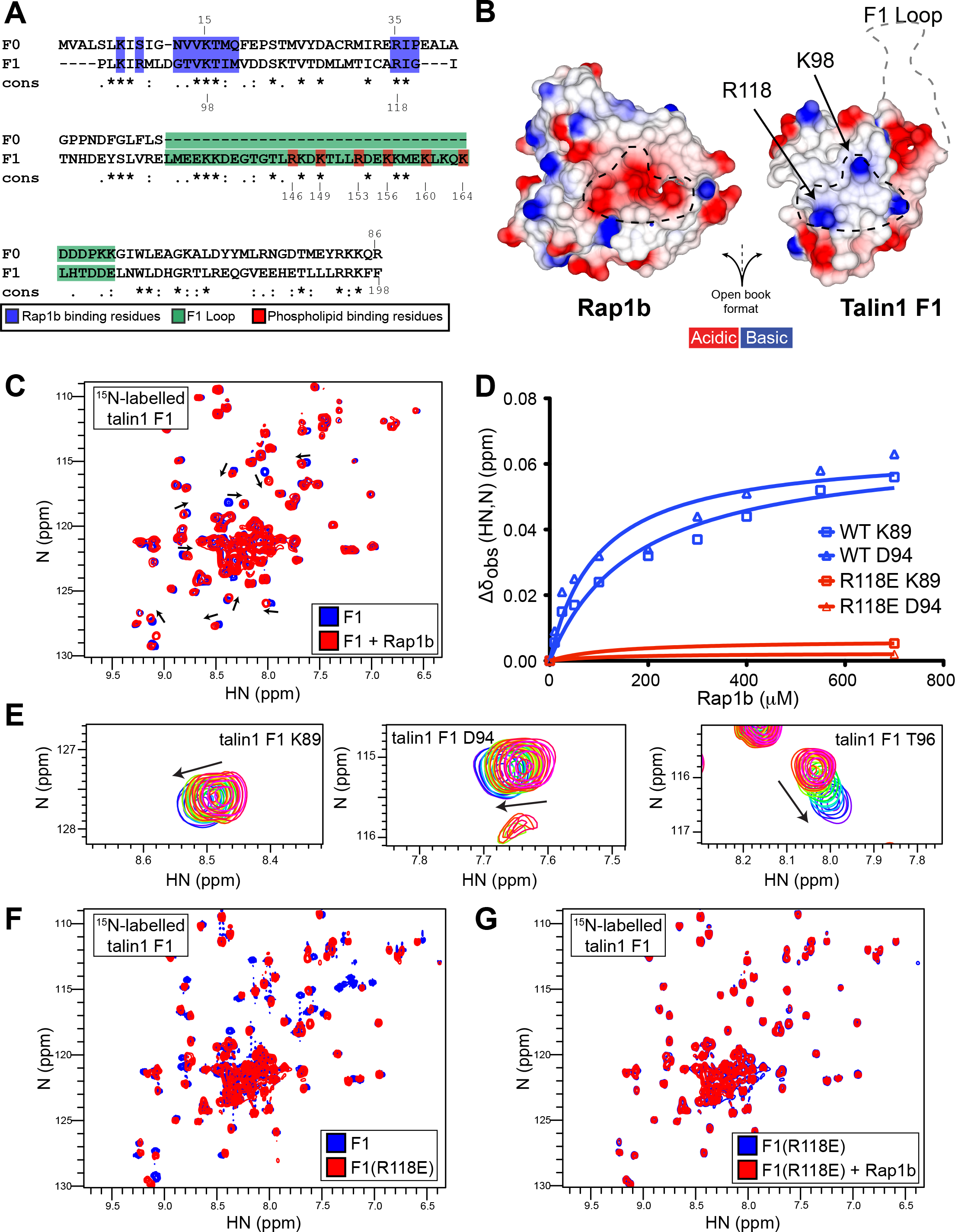
Talin1 F1 subdomain also contains a Rap1b-binding site and the R118E mutation blocks binding to Rap1. **(A)** Amino-acid sequence alignment of the F0 and F1 domains. Invariant residues indicated with “*”, conserved residues with “:”, and semi-conserved residues with “.”. Rap1 binding residues (blue) are >25% buried by formation of the complex. **(B)** Surface electrostatic potential of talin1 F1 (right) and Rap1b (left) binding interface as open book view. The dotted black lines highlight the charge complementarity between the two proteins binding interface. The location of the F1 loop is also shown by a grey dotted line. **(C)** 2D ^1^H,^15^N-sfHMQC spectra of 100 μM ^15^N-labeled talin1 F1 wild-type (blue) and in the presence of 700 μM GMP.PNP Rap1b (red). Specific chemical shift changes are observed indicating binding. **(D)** Representative titration curves for the interaction of Rap1b with talin1 F1 domain wild-type in blue and R118E mutant in red (also see Fig. S2E-H for more examples). The talin1(R118E) mutation has a dramatic effect on Rap1b binding. **(E)** Close up view of the 10-step Rap1b titration for residues K89, D94 and T96. **(F-G)** 2D ^1^H,^15^N-sfHMQC spectra of 100 μM ^15^N-labeled talin1 F1. **(F)** Wild-type (blue) and R118E (red). Almost all peaks are in the same position, suggesting that THD(R118E) is well folded and has a similar fold to wild-type protein. **(G)** Free talin1 F1(R118E) in the absence (blue) and presence of GMP.PNP Rap1b (red) at 1:7 molar ratio. No chemical shift changes are observed at 7-fold excess Rap1b, suggesting that F1(R118E) drastically reduced the affinity of the interaction.

To test the presence of a Rap1 binding site, we prepared ^15^N-labelled talin1 F1 domain for NMR studies. The 2D-sfHMQC of the talin1 F1 domain showed a stable folded protein, with most resonances being highly dispersed (Fig. 2 C, blue). The presence of sharp intense signals located in the center of the spectra is consistent with a 35-residue loop that is unstructured and highly dynamic. Addition of Rap1b caused residue-specific chemical shift changes of talin1 F1 (Fig. 2 C, red), in contrast to the studies of Zhu et al., 2017 using the F1F2 double-domain. As predicted, the non-dispersed resonances from the loop were not affected by Rap1b. Furthermore, we mapped specific chemical shifts, including Arg98 and Arg118 (Fig. S2 A), that clustered to amino acids located in the predicted RA binding interface near the F1 β2 strand (Fig. S2 B). NMR titrations showed concentration-dependent chemical shift perturbations with fast exchange, that suggested a weak affinity of talin1 F1 for Rap1 based on several amino acids located at the binding interface (Fig. 2 D and S2 E-H, blue lines). As highlighted in Fig. 2 E, the chemical shift perturbations were small and made the determination of the affinity less accurate. After careful analysis we found that the apparent Kd values for the F1-Rap1 interaction were varying from low to high μM range as summarized in Fig. S2 C. Analysis of our previous data on the talin1 F0 domain also showed similar variations (Fig. S2 D) (Goult et al., 2010; Lagarrigue et al., 2018), suggesting that this is an intrinsic property of the talin1 F0 and F1 subdomains. Thus the talin1 F1 domain contains an RA domain that can bind Rap1b to a similar extent as the talin1 F0 domain (Goult et al., 2010; Lagarrigue et al., 2018; Zhu et al., 2017).

### Dramatically reduced affinity of talin1(R118E) F1 domain binding to Rap1

Our molecular modeling suggested that talin1 Arg98 and Arg118 play an important role on the surface charge complementarity with Rap1b (Fig. 2 B) and the NMR data showed that their backbone amide chemical shift are affected by binding. We used the acidic Glu mutation because of the predicted stronger effects of charge repulsions between talin1 F1 and the acidic patch on the surface of Rap1b that mediates their interaction. We therefore purified recombinant ^15^N-labelled talin1 F1(R118E) mutant. Comparison of the NMR 2D-sfHMQC of the F1 and F1(R118E) mutant showed only very subtle chemical shift changes (Fig. 2 F); thus, the mutant protein adopts a similar fold to F1. Addition of GMP.PNP Rap1b to F1(R118E) produced no specific chemical shift changes (Fig. 2 G), suggesting that it does not bind to Rap1b. The minimal chemical shift perturbations detected at 7-fold excess of Rap1b with F0(R118E) indicated a substantial reduction in Rap1b affinity (Fig. 2 D and S2 E-H, red lines) as judged by negligible chemical shift for each previously perturbed residue. Thus talin1 F1(R118E) is well folded and has a greatly reduced affinity for Rap1.

### Talin1 F1-Rap1b interaction is important for THD-mediated αIIbβ3 activation in A5 cells

To assess the contribution of the talin1 F0 and F1 interactions with Rap1 on integrin activation, we introduced the mutations R35E, R118E and a double mutant R35E,R118E into the THD (Fig. 3 A) and tested effects on integrin activation in A5 cells. Expression of all THDs induced integrin activation with THD and THD(R35E) producing comparable levels of activation. In sharp contrast, both constructs with F1 mutations, THD(R118E) and THD(R35E,R118E), exhibited a substantially reduced activation (Fig. 3 B, grey bars). Furthermore, we carefully monitored expression of THD-EGFP and its mutants by flow cytometry and observed a strong effect of the F1 mutants at equivalent levels of THD expression on a per cell basis (Fig. 3 C). Immunoblotting confirmed similar level of global expression for all THD mutants (Fig. 3 D). More importantly, co-expression of both Rap1(Q63E) and Rap1GAP1 had no effect on integrin activation when the R118E mutation was introduced in the F1 domain (Fig. 3 B, red and blue bars). Also, THD-induced activation in the presence of Rap1GAP1 was very similar to activation with the THD(R118E) mutant alone, further supporting the conclusion that the Rap1-F1 interaction plays a crucial role in Rap1’s ability to potentiate THD-induced αIIbβ3 activation.

**Figure 3.**
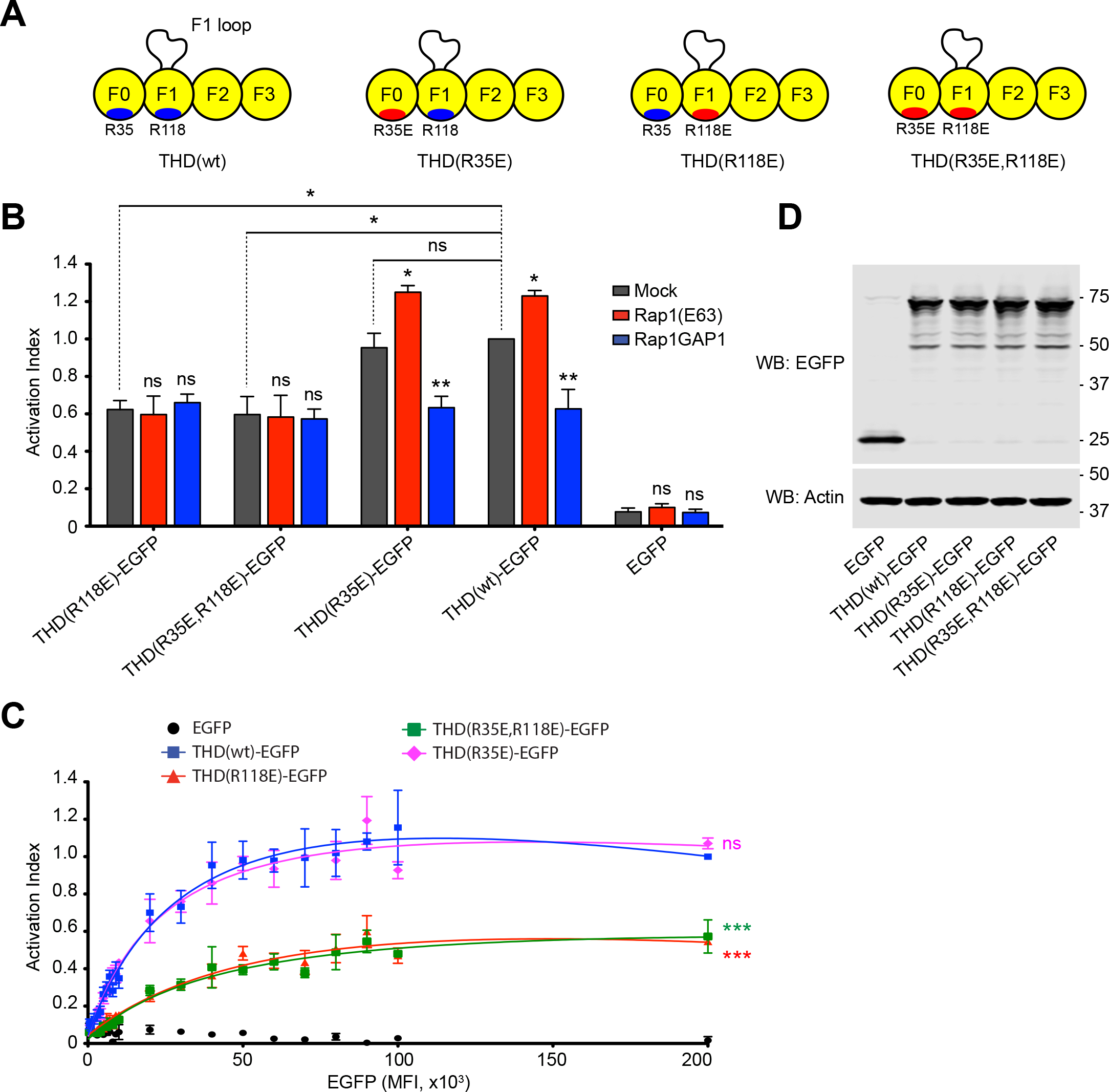
The talin1 F1 domain is responsible for THD Rap1-dependent integrin activation. **(A)** Talin1 head domain (THD) constructs used in panels B-D. **(B-C)** A5 cells stably expressing αIIbβ3 integrin were transfected with cDNA encoding THD-EGFP in combination with Rap1a(Q63E) or Rap1GAP1. Integrin activation was assayed by binding of PAC1 to EGFP positive cells. Transfection of EGFP alone was used as control. **(B)** Bar graphs represent Mean ± SEM of 4 independent experiments, and normalized to THD-EGFP + Mock. Two-way ANOVA with Bonferroni post-test. Each condition was compared to the respective Mock control. ns, not significant; *, p<0.05; **, p<0.01. **(C)** Activation indices were normalized to the maximum value of THD-EGFP and plotted as a function of EGFP MFI. Graphs represent Mean ± SEM of 4 independent experiments. Curve fitting was performed using the total one site-binding model in Prism 5.0 (GraphPad Software). Two-way ANOVA with Bonferroni post-test. Each mutant was compared to THD. ns, not significant; ***, p<0.01. **(D)** Western blot of THD-EGFP expression. Actin was used as a loading control.

### Talin1 F1-loop is important for Rap1 to increase THD-mediated αIIbβ3 activation i

Since both the talin1 F0 and F1 domains contain a Rap1-binding site with similar affinity for Rap1b *in vitro*, and only the THD(R118E) mutant in F1 had an effect on THD-mediated αIIbβ3 activation, we examined differences between the two domains. First, the F1 domain is located adjacent to the F2 domain which contains a critical membrane orientation patch required for activation (Anthis et al., 2009); however, in THD(ΔF1), the F0 is adjacent to F2 yet THD(ΔF1) exhibits little Rap1-dependence (Fig. 1 F). Secondly, the F1 domain contains an unstructured loop implicated in integrin activation. (Goult et al., 2010) showed that this loop has helical propensity and forms a more stable helix on binding to phospholipids. Basic residues, predicted to reside on one surface of the helix, are required for binding to acidic phospholipids (Goult et al., 2010).

To assess the contribution of both the talin1 F1 loop and the interaction with Rap1 on integrin activation, we tested integrin activation by THD wherein the loop was removed, THD(ΔL) (Fig. 4 A). As previously observed (Goult et al., 2010), THD(ΔL) reduced integrin activation (Fig. 4 B, grey bars). Importantly, co-expression of both Rap1(Q63E) and Rap1GAP1 had a negligible effect on integrin activation by THD(ΔL) (Fig. 4 B, red and blue bars). Furthermore, the effect of the double mutant THD(ΔL,R118E) resembled that of THD(ΔL). Immunoblotting confirmed similar level of expression for all THD mutants (Fig. 4 C). 2D-sfHMQC NMR spectra of ^15^N-labelled talin1 F1(ΔL) exhibited highly dispersed resonances consistent with a stable folded protein (Fig. 4 D, blue). Addition of Rap1b caused residues specific chemical shift changes in talin1 F1(ΔL) (Fig. 4 D, red) that mapped to amino acids located in the predicted RA binding interface including Thr96 and Thr99 (Fig. 4 E). Thus, the talin1 F1 loop is important for Rap1-dependence of THD-mediated integrin activation but not for Rap1b binding, indicating that the t loop works in tandem with Rap1 binding in performing this function.

**Figure 4.**
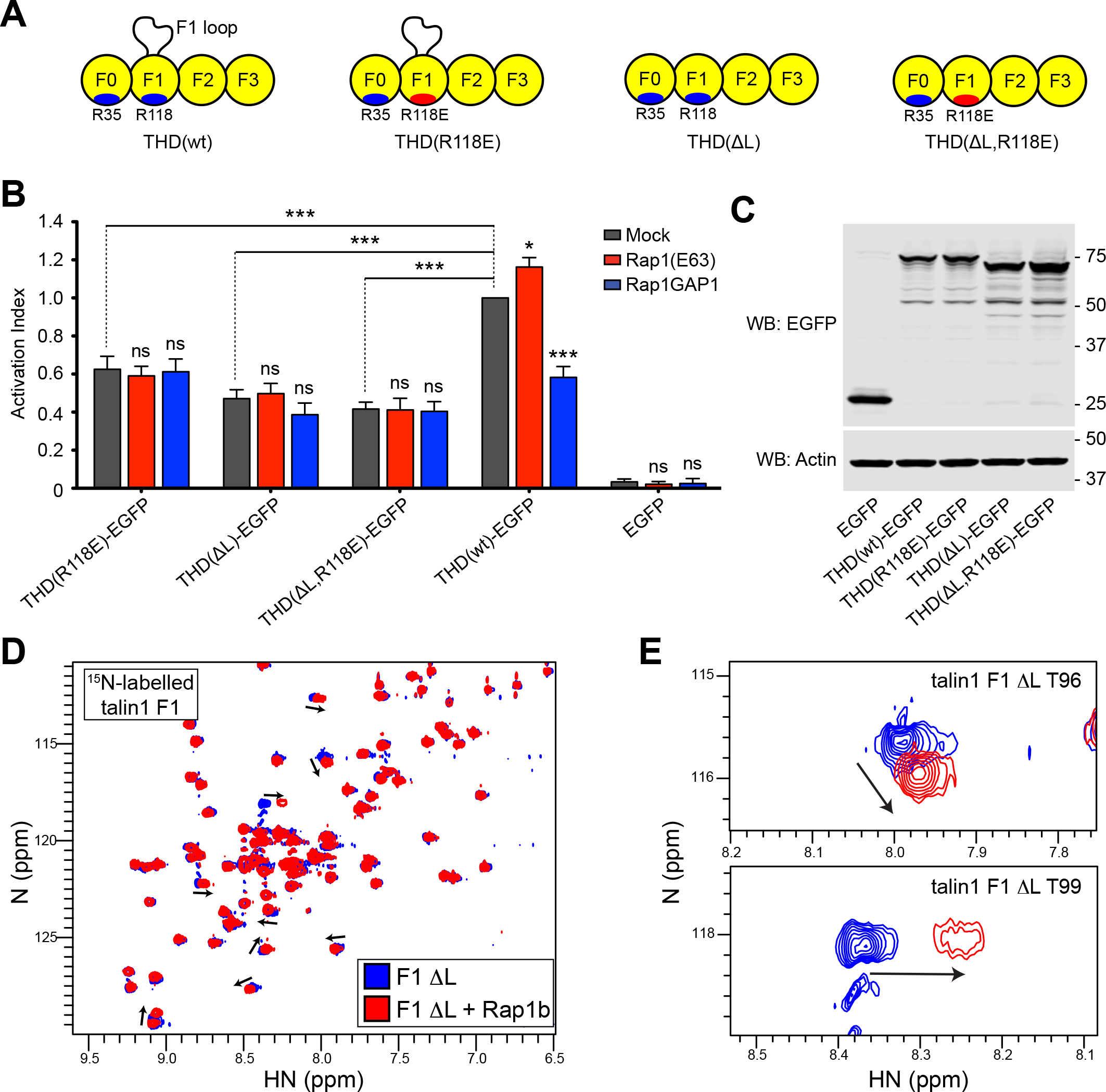
The talin1 F1 domain inserted loop plays a synergistic role with the F1 Rap1 binding site for integrin activation. **(A)** THD constructs used in panels B and C. **(B)** A5 cells stably expressing αIIbβ3 integrin were transfected with cDNA encoding THD-EGFP in combination with Rap1a(Q63E) or Rap1GAP1. Integrin activation was assayed by binding of PAC1 to EGFP positive cells. Transfection of EGFP alone was used as control. Bar graphs represent Mean ± SEM of 4 independent experiments, and normalized to THD-EGFP + Mock. Two-way ANOVA with Bonferroni post-test. Each condition was compared to the respective Mock control. ns, not significant; *, p<0.05; ***, p<0.001. **(C)** Western blot of THD-EGFP expression. Actin was used as a loading control. **(D)** 2D ^1^H,^15^N-sfHMQC spectra of 100 μM ^15^N-labeled talin1 F1 ΔL (blue) and in the presence of 225 μM GMP.PNP Rap1b (red). Specific chemical shift changes are observed indicating binding. **(E)** Close up view of the chemical shift changes for residues T96 and T99.

To further validate the proposed formation of a helix mechanism whereby the talin1 F1 loop plays a role in integrin activation, we introduced three charge reversal mutations in basic residues on the surface of the predicted helix (R146E,R153E,K156E (3EL)) to create a charge repulsion with the negatively charged PI(4,5)P_2_ of the plasma membrane (Fig. S3 A) that blocks lipid binding (Goult et al., 2010). THD(3EL) was even less active than THD(ΔL) in activating αIIbβ3 (Fig. S3 B). Immunoblotting confirmed similar level of expression for all THD mutants (Fig. S3 C).

Since we used the THD in our experiments, we also wanted to test the effect of the F1 mutants in full-length talin1. To ensure maximal Rap1 activation we expressed Rap1(Q63E) in A5 cells and noticed no increase in αIIbβ3 activation consistent with the modest levels of talin in these cells (Fig. S3 D), as previously reported (Han et al., 2006). Co-expression of talin1 increased activation as did the talin1(R35E) mutant. Importantly, both the talin1(R118E) and talin1(ΔL) reduced integrin activation to a similar extent. Indeed, the effect of the combined talin1(ΔL,R35,R118E) mutant was similar to talin1(R118E) alone, emphasizing: i) the critical role of Rap1 binding F1; and ii) the Rap1-dependence of talin1-induced activation depends on the loop in F1 thus explaining why F1 is more important than F0 in this function.

### Rap1 binding to THD F0 and F1 facilitates its recruitment to paxillin-containing adhesions

To further evaluate the role of Rap1-talin1 interaction in integrin functions, we examined the recruitment of THD to adhesion structures in adherent 3T3 fibroblasts. THD localized to paxillin-positive adhesions (Fig. 5 A). A previous report revealed the role of Rap1 binding to the F0 domain in talin1 recruitment to adhesion sites (Zhu et al., 2017). Similarly, we found that co-localization of THD(R35E) mutant with paxillin was reduced (Fig. 5 A-B). Remarkably, THD(R118E) mutant showed an even more impaired recruitment to paxillin-positive adhesions (Fig. 5 A-B) resulting in more diffuse cytoplasmic localization. Thus, Rap1 binding to each Rap1 binding site in THD is important for recruitment to paxillin-containing adhesions.

**Figure 5.**
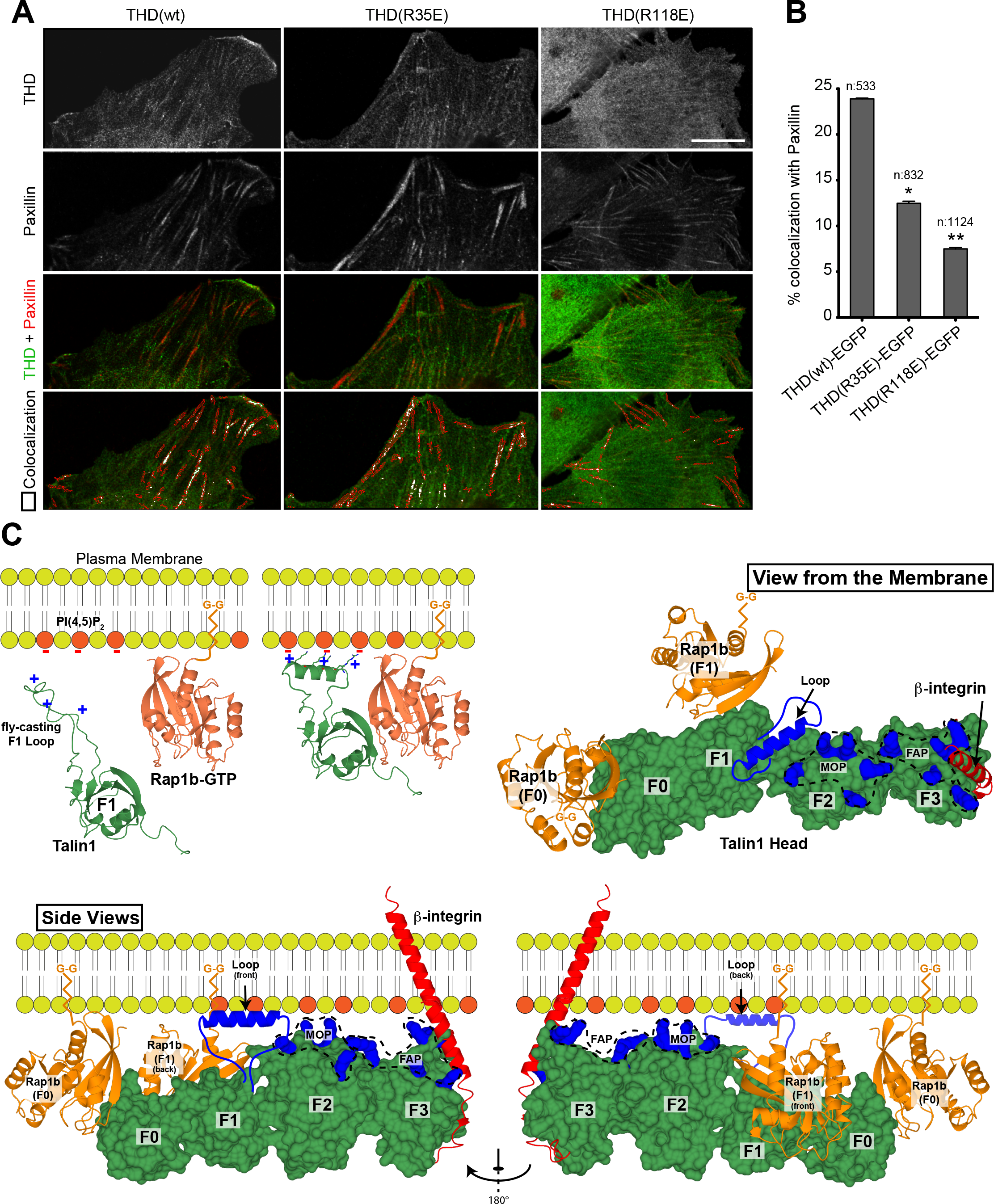
The role of talin1 F1 domain talin recruitment. **(A)** Airyscan LSM Image MIP projections of 3T3 fibroblasts expressing both THD-EGFP (green) and mRby2-Paxillin (red). Bottom panel shows the merged image and the paxillin rich ROIs (red lines) with quantitative colocalization of both THD to paxillin in peripherally labelled paxillin rich focal adhesions. The images clearly show a diminishment of co-localized signal of THD to paxillin between the WT control and THD(R35E) mutant, which is further decreased in the THD(R118E) mutant. Quantitation results are displayed in **(B)**. The number of analyzed adhesions is indicated for each condition. **(C)** A model of talin1 F1 domain in the cytosol with a positively charged unstructured loop (green) and membrane-associated Rap1b-GTP by its C-terminal geranyl-geranyl (G-G) moiety (orange). Negatively charged phospholipids PI(4,5)P_2_ are colored in red. On proximity of the plasma membrane, the low-affinity talin1 F1 Ras-associating (RA) domain probes for Rap1 and the F1 loop seeks negatively charged phospholipids. On contact with Rap1 and negatively charged phospholipids, the F1 RA domain interacts with Rap1 and the F1 loop helical state is favored resulting in cluster of positive charges on one side of the helix. View of THD (green) as seen from the membrane is displayed in the top right panel. Bottom panel shows view of THD as seen from the side. The views highlight the position of the THD F0, F1, F2 and F3 subdomains and the Rap1b (orange) bound to the F0 and F1 subdomains. Both Rap1b C-terminal geranyl-geranyl (G-G) moieties are pointing towards the membrane, as for the F1 loop, the F2 membrane orientation patch (MOP), the F3 association patch (FAP), and the position of the F3 β-integrin. The regions known to interact with the negatively charged phospholipids are shown in blue; the F1 fly-casting loop is shown as a helix, the F2 membrane orientation patch (MOP), and the F3 association patch (FAP). The β-integrin transmembrane and cytoplasmic tail is shown in red bound to the F3 subdomain. Two Rap1b (orange) molecules are shown with their C-terminal geranyl-geranyl (G-G) moiety inserted in the membrane. One Rap1b is bound to the F0 and one to the F1 subdomain. This model summarize the multiple known interactions of THD at the plasma membrane; the negatively charged PI(4,5)P_2_, Rap1 and integrin β-tails.

The studies reported here provide new insight into how talin1 activates integrins, a process central to mammalian development and numerous physiological functions. Studies in A5 cells led to the conclusion that talin1 is required for integrin activation, established the importance of specific structural elements in the integrin and in THD, and identified the importance of lipid binding residues (Anthis et al., 2009; Goult et al., 2010; Tadokoro et al., 2003; Wegener et al., 2007). The A5 cells were also instrumental in studies that showed how Rap1, by engaging RIAM enabled RIAM to recruit talin1 to integrins (Han et al., 2006; Lee et al., 2009); a key pathway in leukocyte trafficking and formation of the immune synapse (Lagarrigue et al., 2017). Nevertheless, Rap1 can activate integrins in the absence of RIAM or of obvious candidate effectors. Goult’s pioneering studies suggested that talin1 F0 could enable talin1 to itself be an effector (Goult et al., 2010), an idea further advanced by Zhu et al. 2017; however we found that the F0 domain made a minor contribution in mammals. We now find that the F1 domain contains a second Rap1 binding site and that this site functions in conjunction with a unique inserted loop to enable Rap1 to regulate activation. The proximity of the putative membrane binding helix of the loop to the geranyl-geranyl moiety of Rap1 bound to F1 provides a cogent model to explain the complementary role of these two membrane-binding sites in integrin activation (Fig. 5 C). Indeed, the now more complete picture of how THD interacts with the membrane shows that an extended series of membrane binding sites serve to stabilize the weak interaction of talin1 F3 with the integrin β cytoplasmic domain to explain the membrane dependence of talin1-induced activation (Ye et al., 2010). Our studies show how Rap can add to those membrane-binding sites to enable activation.

Our studies raise several new questions. In mice, Rap1 binding to talin F0 is clearly dispensable for normal development and has only a modest effect on activation of αIIbβ3 and platelet function. In stark contrast, in Dictyostelium (Plak et al., 2016), and Drosophila (Camp et al., 2018) blocking Rap1 binding to talin F0 has severe consequences for morphogenesis. We note that the Dictyostelium TalinB F1 has no loop and in Drosophila talin the loop sequence is not conserved, suggesting that evolution has endowed mammalian F1 with a loop capable of supporting most Rap1-dependent integrin function, even in the absence of F0’s contribution. Absent this enhanced function of F1, F0’s contribution may become indispensable. This hypothesis will be testable by future reverse genetic studies in which the developmental and physiological effects of Rap1 binding-defective F1 mutations can be assessed alone or in combination with Rap1 binding-defective F0 mutations. The future reverse genetic experiments will also enable studies to assess the importance of Rap1 binding to talin in cells, such as leukocytes, that possess sufficient MRL proteins (e.g. RIAM) to provide an alternative mechanism by which a Rap1 effector can engage talin, leading to its interaction with and activation of integrins. Finally, our findings will enable future studies to characterize the role of each of these Rap1 binding sites in activation in cells in which talin1 and Rap1 are in varying abundance and to assess their roles in stability of integrin-mediated adhesions and resulting mechanotransduction.

## Materials and methods

### Integrin activation in CHO cells

Cells were cultured in DMEM (Corning) supplemented with 10% fetal bovine serum (Sigma-Aldrich), and 100 U/ml penicillin and 100 μg/ml streptomycin (Gibco). The sequences encoding murine talin1 head domain (THD, aa 1-433) or full-length talin1 were cloned into pEGFP-N1 (Clontech). The sequences encoding human Rap1a(Q63E) and Rap1GAP1 were cloned into p3xFlag7.1(-) (Clontech). Transient transfection was performed using TransIT-LT1 Reagent (Mirus) according to the manufacturer’s protocol. PAC-1 binding assay in A5 cellls was conducted as previously described (Frojmovic et al., 1991). Briefly, cells were harvested by using trypsin one day after transfection and washed once in HBSS buffer (containing calcium and magnesium, Gibco) supplemented with 1% (w:v) bovine serum albumin (Sigma-Aldrich). PAC-1 IgM (ascites, 1:200) was incubated with cells for 30 min at room temperature prior to staining with the appropriate Alexa Fluor 647 secondary antibody (Life Technologies) for 30 min on ice. Cells were analyzed by flow cytometry using a FACS Accuri C6 Plus (BD Biosciences) and gated on EGFP-positive events. Integrin activation was defined as αIIbβ3 specific ligand binding corrected for α_IIb_β_3_ expression, and was calculated as 100×(MFI_i_-MFI_0_)/ΔMFI_D57_ (where MFI_i_ = mean fluorescence intensity of bound PAC-1; MFI_0_ = mean fluorescence intensity of bound PAC-1 in the presence of 10 μM Eptifibatide; and ΔMFI_D57_ = specific fluorescence intensity of anti-α_IIb_β_3_ D57 antibody). PAC1 (Shattil et al., 1985) and D57 (O’Toole et al., 1994) antibodies have been previously described.

### Protein expression and purification

The murine talin1 residues 1-400 (THD), 84-400 (F1-F3), 84-196 (F1), and 84-196 Δ141-170 (F1 Δloop) were cloned into the expression vector pETM-11 (His-tagged, EMBL) and expressed in Escherichia coli BL21 Star (DE3) cultured in minimal medium for ^15^N-labeled samples for NMR or LB for unlabeled protein. Briefly, recombinant His-tagged proteins was purified by nickel-affinity chromatography, the His-tag removed by cleavage with Tobacco Etch Virus protease overnight, and the protein further purified by size exclusion chromatography using a Superdex-75 (26/600) column (GE Healthcare). The column was pre-equilibrated and run with (NMR-buffer) 20 mM Sodium Phosphate, 50 mM NaCl, 3 mM MgCl_2_, 2 mM DTT, pH 6.5. Human Rap1 isoform Rap1b (residues 1– 167) cloned into pTAC vector in the *E. coli* strain CK600K was the generous gift of Professor Alfred Wittinghofer (Max Planck Institute of Molecular Physiology, Dortmund, Germany). Untagged Rap1b was purified by ion exchange, followed by Superdex-75 (26/600) gel filtration as previously described (Gingras et al., 2016). The column was pre-equilibrated and run with NMR-buffer.

### NMR spectroscopy

NMR samples of all the protein constructs were prepared in NMR-buffer containing 0.1 mM GMP.PNP and 5% (v/v) of ^2^H_2_O. All 2D [^1^H,^15^N]-sfHMQC spectra were recorded at 298K. To test THD binding to Rap1b, 100 μM ^15^N-labelled GMP.PNP-bound Rap1b was incubated with 250 μM unlabelled THD or THD(ΔF0). Titration curves for the interaction of talin1 F1 with GMP.PNP-bound Rap1b were determined using 100 μM ^15^N-labelled F1 in NMR-buffer. Chemical shift changes (Δδ_obs (HN,N)_) were calculated using CcpNmr Analysis, “follow shift changes” function and analyzed with the one site binding model to determine the apparent Kd value in Prism 5.0 (GraphPad Software) (Vranken et al., 2005).

### Western blotting

Cells were lysed in Laemmli sample buffer. Lysates were subjected to a 4-20% gradient SDS-PAGE. Polyclonal serum directed against EGFP was raised in rabbit (Abgent). Antibody against β-actin (AC-15) was from Sigma-Aldrich. The appropriate IRDye/Alexa Fluor-coupled secondary antibodies were from LI-COR. Nitrocellulose membranes were scanned using an Odyssey CLx infrared imaging system (LI-COR) and blots were processed using Image Studio Lite software (LI-COR).

### Microscopy

Flp-In™ 3T3 cells (Life technologies) were cultured in DMEM (Corning) supplemented with 10% fetal bovine serum (Sigma-Aldrich), and 100 U/ml penicillin and 100 μg/ml streptomycin (Gibco). The sequence encoding mRuby2 fused to the C-terminus of Paxillin was cloned into the pMSCV retroviral backbone (Addgene). Transduced mRuby2-positive cells were single cell cloned by automated cell deposition unit using a BD FACSAria (BD biosciences). A selected clone was then transfected with plasmids encoding EGFP fused to the C-terminus of murine THD (aa 1-433). THD-EGFP wild-type or mutant sequences downstream of the Tet-On responsive element were cloned into the pcDNA5/FRT plasmid (Life Technologies) expressing the Tet-On transactivator from the mouse PGK promoter. Cells were co-transfected with pcDNA5 constructs and the pOG44 Flp recombinase expression vector using TransIT-LT1 (Mirus). Transformants were selected in presence of 200 μg/ml of Hygromycin B (Life Technologies). THD-EGFP expression was induced by adding 100 ng/ml of doxycycline (Sigma-Aldrich) overnight. Cells were trypsinized and seeded on top of glass coverslips coated with 10 μg/ml of human fibrinogen (Sigma-Aldrich). Cells were fixed two hours later with 2% formaldehyde (Tousimis) for 20 min at 37°C, permeabilized with Triton X-100, and stained with DAPI and Phalloidin conjugated to Alexa Fluor 647 (Life Technologies). Three dimensional stacks of images were acquired using a Zeiss Laser Scanning Confocal Microscope LSM Airy Scan 880, equipped with a 63x (1.46na) objective (using a 0.17 micron step interval) and an automated piezo stage. Each image consisted of z-stacks of multiple frames that were first processed from a raw Airyscan image to a final integrated, corrected and deconvolved image that was then flattened as a maximum intensity projection using the ZEN software (Zeiss Inc.). Only the optical two image slices that were closest to the coverslip were projected in order to analyze localized proteins that are on the basal surface of the cell: peripheral paxillin rich adhesions. The Images were then further processed in Image Pro Premier 10 [IPP10] (Media Cybernetics) in the following manner. MIP images were calibrated and auto-traced/masked for paxillin rich signals using a smart segmentation macro inherent in IPP10 that, based on control images, was able to define all relevant paxillin rich adhesions in the cells and outline them. These masked outlines were used as regions of interest (ROIs) from which a colocalization module in IPP10 auto calculated and displayed on the image (Pseudo-colored in white) the amount of colocalization between paxiilin rich zones (red) and THD rich zones (green) as seen in last panel of Figure 5A. The relevant fluorescent signal range in the colocalization module was between (60-256 levels of grey), well above background controls. All colocalization values from IPP10 were exported directly into Excel where they were further processed. The experiment was repeated twice examining on average 800 paxillin rich adhesions over an average of 20 cells per category.

### Reagents

GMP.PNP was purchased from Sigma.

### Statistical analysis

Statistical significance was assayed by a two-tailed t-test for single comparisons or ANOVA for multiple comparisons with a Bonferroni post hoc test. A p value <0.05 was considered significant.

## Online supplemental material

Fig. S1 shows A5 cells stably expressing αIIbβ3 integrin transfected with cDNA encoding THD(wt)-EGFP or THD(TM)-EGFP, or in the presence of MnCl_2_. Fig. S2 summarizes the effects of Rap1 binding on the talin F1 subdomain by NMR spectroscopy. Fig. S3 shows the effects of transfecting THD F1 loop mutants and full-length talin1 mutants in A5 cells.

## Acknowledgments

This work was supported by the National Institutes of Health - National Heart, Lung and Blood Institute grants HL 139947 (M.H.G.), K01HL 133530-01 (M.A.L.R), and HL 078784 (M.H.G. and K.L.). American Heart Association Career Development Award 18CDA34110228 (F.L.) and Grant-In-Aid 16GRNT29650005 (A.R.G.); and Zeiss LSM880 was supported by NIH S10OD021831. The authors declare no competing financial interests.

## Author contributions

Contribution: A.R. Gingras, F. Lagarrigue and M.H. Ginsberg conceived the study, designed experiments, interpreted data, and wrote the manuscript. Alexandre R. Gingras, Frederic Lagarrigue, Monica N. Cuevas, Marcus Zorovich, Andrew J. Valadez, Wilma McLaughlin, William Kiosses and Nicolas Seban performed and analyzed experiments. Miguel Alejandro Lopez-Ramirez and Klaus Ley provided vital reagents and critical expertise.

## Abbreviations

Abbreviations used in this paper:

THD: talin1 head domain

CHO cells: Chinese hamster ovary

---

**Supplementary Figure 1.**
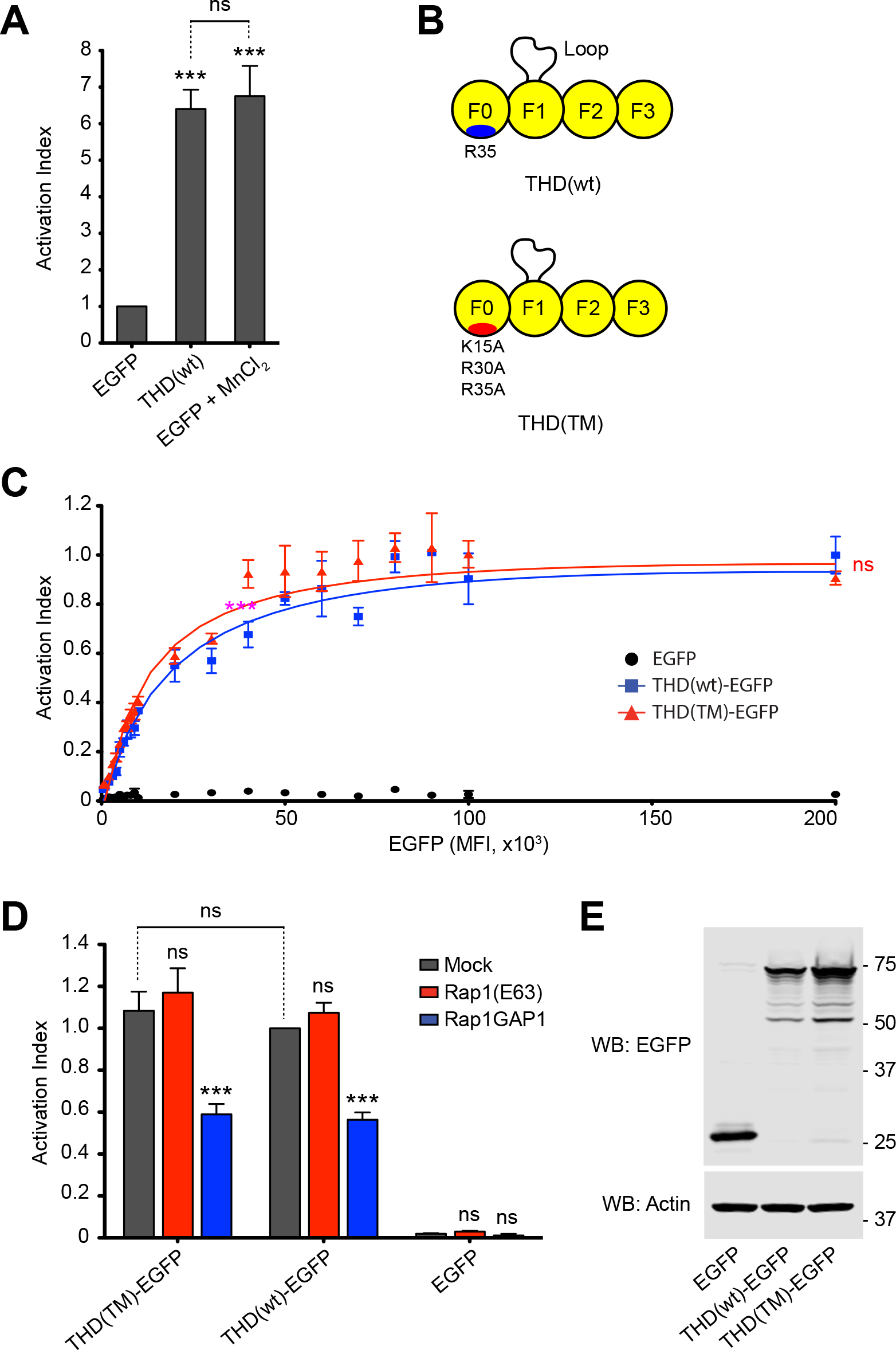
THD(wt)-EGFP, THD(TM)-EGFP and MnCl_2_ show similar levels of integrin activation. **(A)** THD and Mn++ activate integrin αIIbβ3 to a similar extent. A5 cells stably expressing αIIbβ3 integrin were transfected with cDNA encoding EGFP or THD-EGFP. Integrin activation was assayed by binding of PAC1 to EGFP positive cells in presence or absence of 1 mM MnCl_2_. Bar graphs represent Mean ± SEM of 3 independent experiments, and normalized to the EGFP condition. Two-way ANOVA with Bonferroni post-test. Each condition was compared to the EGFP control. ***, p<0.001. Comparisons to the THD condition is displayed in green. ns, not significant. **(B-E)** Talin1(K15A,R30A,R35A) (TM) triple mutation does not impede αIIbβ3 integrin activation in CHO cells. **(B)** THD constructs used in panels C and D. **(C-D)** A5 cells stably expressing αIIbβ3 integrin were transfected with cDNA encoding THD-EGFP alone (C) or in combination with Rap1a(Q63E) or Rap1GAP1 (D). Integrin activation was assayed by binding of PAC1 to EGFP positive cells. Transfection of EGFP alone was used as control. **(C)** Activation indices were normalized to the maximum value of THD-EGFP and plotted as a function of EGFP-MFI. Graphs represent Mean ± SEM of 3 independent experiments. Curve fitting was performed using the total one site-binding model in Prism 5.0 (GraphPad Software). Two-way ANOVA with Bonferroni post-test. Each mutant was compared to THD. ns, not significant. **(D)** Bar graphs represent Mean ± SEM of 3 independent experiments, and normalized to THD-EGFP + Mock. Two-way ANOVA with Bonferroni post-test. Each condition was compared to the respective Mock control. ns, not significant; ***, p<0.001. No significant differences between wild-type and the triple mutant (TM) were detected. **(E)** Expression of the THD(TM)-EGFP triple mutant was assayed by Western blotting.

**Supplementary Figure 2.**
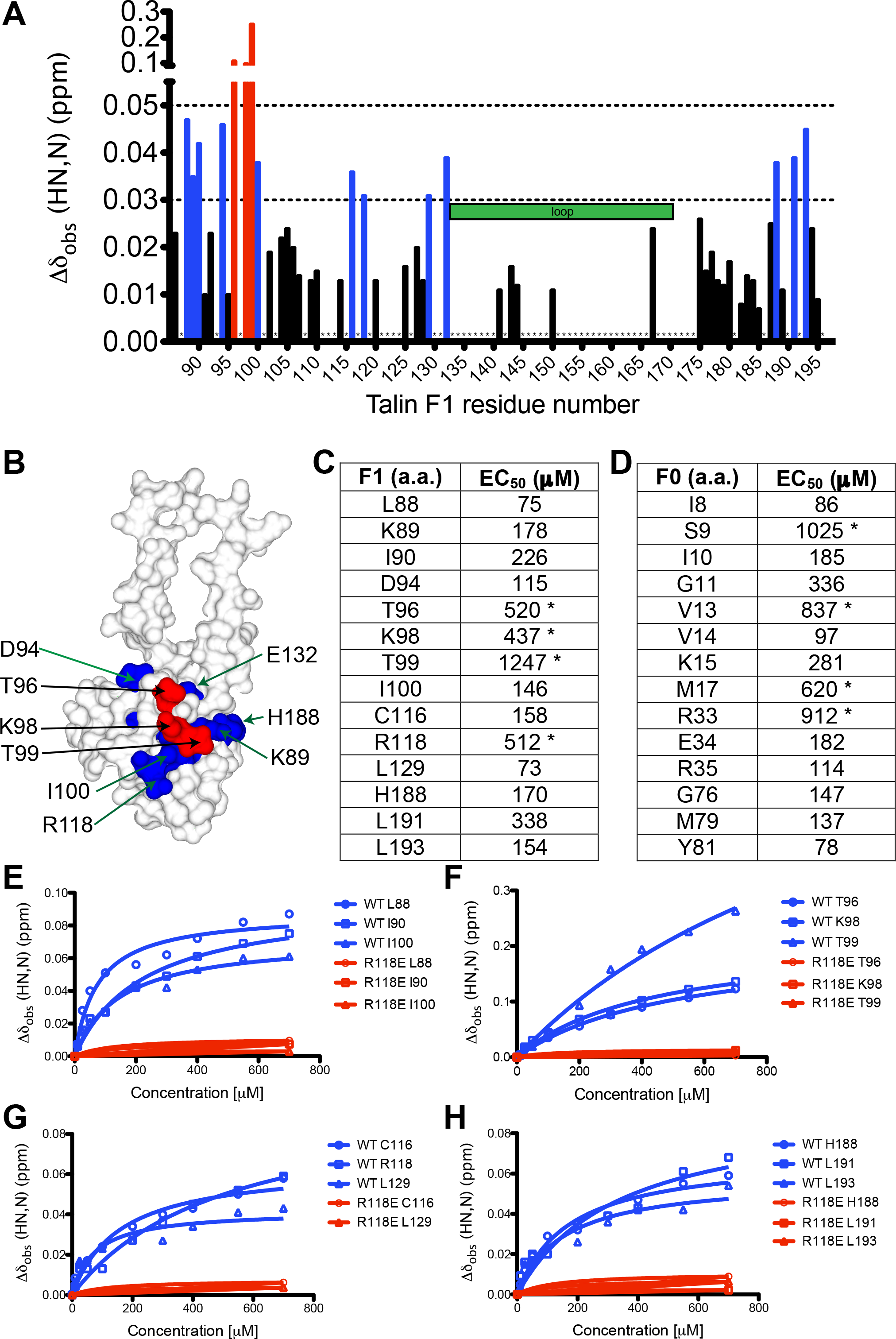
Characterization of the Rap1 binding to talin F1 subdomain by NMR. **(A)** Summary of chemical shift changes as a plot against the residue number of talin1 F1 domain for assigned residues with high confidence using the previously assigned spectra from (Goult et al., 2010). **(B)** Mapping of the significant changes on the F1 structure with the same color code as panel A. **(C-D)** The calculated apparent Kd values using amino acids backbone amide chemical shift perturbations for the talin1: (C) F1; and (D) F0 domains. The apparent Kd values are observed in similar μM ranges, suggesting similar affinities F1 and F0 for Rap1b. The asterisks highlight the larger values. **(E-H)** Titration curves for the interaction of talin1 F1 with Rap1b for residues assigned with confidence using the previously published assignment (Goult et al., 2010).

**Supplementary Figure 3.**
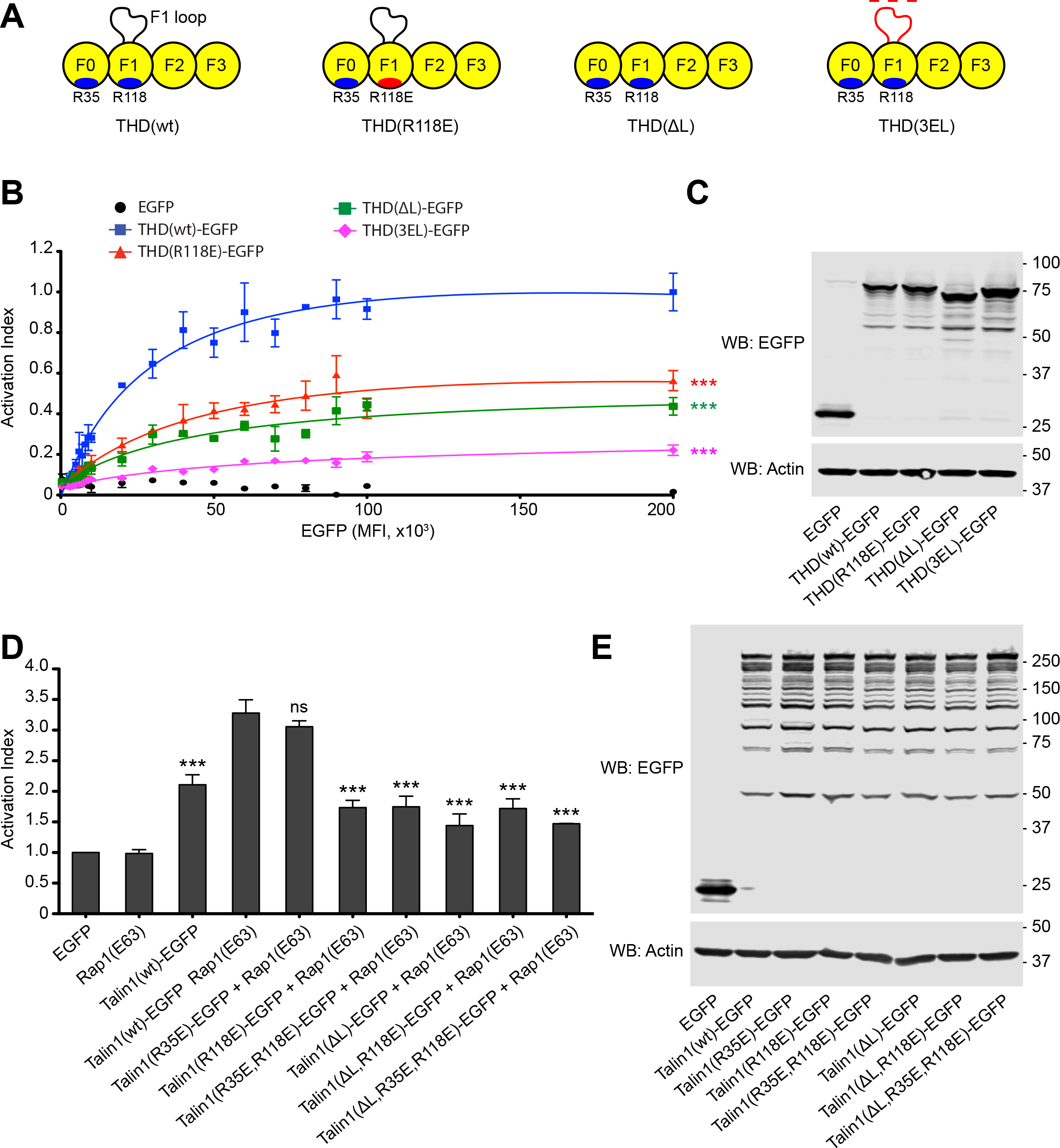
The role of talin1 basic residues of the F1 loop and full-length talin1 mutants in integrin activation. **(A)** THD constructs used in panels B and C. **(B)** A5 cells stably expressing αIIbβ3 integrin were transfected with cDNA encoding THD-EGFP. Integrin activation was assayed by binding of PAC1 to EGFP positive cells. Transfection of EGFP alone was used as control. Activation indices were normalized to the maximum value of THD-EGFP and plotted as a function of EGFP MFI. Graphs represent Mean ± SEM of 3 independent experiments. Curve fitting was performed using the total one site-binding model in Prism 5.0 (GraphPad Software). Two-way ANOVA with Bonferroni post-test. Each mutant was compared to THD. ***, p<0.01. **(C)** Western blot of THD-EGFP expression. Actin was used as a loading control. **(D)** A5 cells stably expressing αIIbβ3 integrin were transfected with cDNA encoding full-length talin1 fused to EGFP in combination with Rap1a(Q63E). Integrin activation was assayed by binding of PAC1 to EGFP positive cells. Bar graphs represent Mean ± SEM of 3 independent experiments, and normalized to talin1(wt)-EGFP alone. One-way ANOVA with Bonferroni post-test. Each condition was compared to the talin1(wt)-EGFP alone control. ns, not significant; ***, p<0.001. **(E)** Western blot of talin1-EGFP expression. Actin was used as a loading control.

## References

Anthis, N.J., K.L. Wegener, F. Ye, C. Kim, B.T. Goult, E.D. Lowe, I. Vakonakis, N. Bate, D.R. Critchley, M.H. Ginsberg, and I.D. Campbell. 2009. The structure of an integrin/talin complex reveals the basis of inside-out signal transduction. The EMBO journal. 28:3623–3632.

Bromberger, T., S. Klapproth, I. Rohwedder, L. Zhu, L. Mittmann, C.A. Reichel, M. Sperandio, J. Qin, and M. Moser. 2018. Direct Rap1/Talin1 interaction regulates platelet and neutrophil integrin activity in mice. Blood. 132:2754–2762.

Calderwood, D.A., I.D. Campbell, and D.R. Critchley. 2013. Talins and kindlins: partners in integrin-mediated adhesion. Nature reviews. Molecular cell biology. 14:503–517.

Camp, D., A. Haage, V. Solianova, W.M. Castle, Q.A. Xu, E. Lostchuck, B.T. Goult, and G. Tanentzapf. 2018. Direct binding of Talin to Rap1 is required for cell-ECM adhesion in Drosophila. Journal of cell science. 131.

Critchley, D.R., and A.R. Gingras. 2008. Talin at a glance. Journal of cell science. 121:1345–1347.

Frojmovic, M.M., T.E. O’Toole, E.F. Plow, J.C. Loftus, and M.H. Ginsberg. 1991. Platelet glycoprotein IIb-IIIa (alpha IIb beta 3 integrin) confers fibrinogen- and activation-dependent aggregation on heterologous cells. Blood. 78:369–376.

Gingras, A.R., W. Puzon-McLaughlin, A.A. Bobkov, and M.H. Ginsberg. 2016. Structural Basis of Dimeric Rasip1 RA Domain Recognition of the Ras Subfamily of GTP-Binding Proteins. Structure. 24:2152–2162.

Goult, B.T., M. Bouaouina, P.R. Elliott, N. Bate, B. Patel, A.R. Gingras, J.G. Grossmann,G.C. Roberts, D.A. Calderwood, D.R. Critchley, and I.L. Barsukov. 2010. Structure of a double ubiquitin-like domain in the talin head: a role in integrin activation. The EMBO journal. 29:1069–1080.

Haling, J.R., S.J. Monkley, D.R. Critchley, and B.G. Petrich. 2011. Talin-dependent integrin activation is required for fibrin clot retraction by platelets. Blood. 117:1719–1722.

Han, J., C.J. Lim, N. Watanabe, A. Soriani, B. Ratnikov, D.A. Calderwood, W. Puzon-McLaughlin, E.M. Lafuente, V.A. Boussiotis, S.J. Shattil, and M.H. Ginsberg. 2006. Reconstructing and deconstructing agonist-induced activation of integrin alphaIIbbeta3. Current biology : CB. 16:1796–1806.

Hynes, R.O. 2002. Integrins: bidirectional, allosteric signaling machines. Cell. 110:673–687.

Klapproth, S., M. Sperandio, E.M. Pinheiro, M. Prunster, O. Soehnlein, F.B. Gertler, R. Fassler, and M. Moser. 2015. Loss of the Rap-1 effector RIAM results in leukocyte adhesion deficiency due to impaired beta2 integrin function in mice. Blood.

Lagarrigue, F., F.B. Gertler, M.H. Ginsberg, and J.M. Cantor. 2017. Cutting Edge: Loss of T Cell RIAM Precludes Conjugate Formation with APC and Prevents Immune-Mediated Diabetes. Journal of immunology. 198:3410–3415.

Lagarrigue, F., A.R. Gingras, D.S. Paul, A.J. Valadez, M.N. Cuevas, H. Sun, M.A. Lopez-Ramirez, B.T. Goult, S.J. Shattil, W. Bergmeier, and M.H. Ginsberg. 2018. Rap1 binding to the talin 1 F0 domain makes a minimal contribution to murine platelet GPIIb-IIIa activation. Blood advances. 2:2358–2368.

Lee, H.S., C.J. Lim, W. Puzon-McLaughlin, S.J. Shattil, and M.H. Ginsberg. 2009. RIAM activates integrins by linking talin to ras GTPase membrane-targeting sequences. The Journal of biological chemistry. 284:5119–5127.

Nieswandt, B., M. Moser, I. Pleines, D. Varga-Szabo, S. Monkley, D. Critchley, and R. Fassler. 2007. Loss of talin1 in platelets abrogates integrin activation, platelet aggregation, and thrombus formation in vitro and in vivo. The Journal of experimental medicine. 204:3113–3118.

O’Toole, T.E., Y. Katagiri, R.J. Faull, K. Peter, R. Tamura, V. Quaranta, J.C. Loftus, S.J. Shattil, and M.H. Ginsberg. 1994. Integrin cytoplasmic domains mediate inside-out signal transduction. The Journal of cell biology. 124:1047–1059.

Petrich, B.G., P. Fogelstrand, A.W. Partridge, N. Yousefi, A.J. Ablooglu, S.J. Shattil, and M.H. Ginsberg. 2007a. The antithrombotic potential of selective blockade of talin-dependent integrin alpha IIb beta 3 (platelet GPIIb-IIIa) activation. The Journal of clinical investigation. 117:2250–2259.

Petrich, B.G., P. Marchese, Z.M. Ruggeri, S. Spiess, R.A. Weichert, F. Ye, R. Tiedt, R.C. Skoda, S.J. Monkley, D.R. Critchley, and M.H. Ginsberg. 2007b. Talin is required for integrin-mediated platelet function in hemostasis and thrombosis. The Journal of experimental medicine. 204:3103–3111.

Plak, K., H. Pots, P.J. Van Haastert, and A. Kortholt. 2016. Direct Interaction between TalinB and Rap1 is necessary for adhesion of Dictyostelium cells. BMC cell biology. 17:1.

Shattil, S.J., J.A. Hoxie, M. Cunningham, and L.F. Brass. 1985. Changes in the platelet membrane glycoprotein IIb.IIIa complex during platelet activation. The Journal of biological chemistry. 260:11107–11114.

Shattil, S.J., C. Kim, and M.H. Ginsberg. 2010. The final steps of integrin activation: the end game. Nature reviews. Molecular cell biology. 11:288–300.

Simonson, W.T., S.J. Franco, and A. Huttenlocher. 2006. Talin1 regulates TCR-mediated LFA-1 function. Journal of immunology. 177:7707–7714.

Song, X., J. Yang, J. Hirbawi, S. Ye, H.D. Perera, E. Goksoy, P. Dwivedi, E.F. Plow, R. Zhang, and J. Qin. 2012. A novel membrane-dependent on/off switch mechanism of talin FERM domain at sites of cell adhesion. Cell research. 22:1533–1545.

Stefanini, L., R.H. Lee, D.S. Paul, E.C. O’Shaughnessy, D. Ghalloussi, C.I. Jones, Y. Boulaftali, K.O. Poe, R. Piatt, D.O. Kechele, K.M. Caron, K.M. Hahn, J.M. Gibbins, and W. Bergmeier. 2018. Functional redundancy between RAP1 isoforms in murine platelet production and function. Blood.

Stritt, S., K. Wolf, V. Lorenz, T. Vogtle, S. Gupta, M.R. Bosl, and B. Nieswandt. 2015. Rap1-GTP-interacting adaptor molecule (RIAM) is dispensable for platelet integrin activation and function in mice. Blood. 125:219–222.

Su, W., J. Wynne, E.M. Pinheiro, M. Strazza, A. Mor, E. Montenont, J. Berger, D.S. Paul, W. Bergmeier, F.B. Gertler, and M.R. Philips. 2015. Rap1 and its effector riam are required for lymphocyte trafficking. Blood.

Sun, H., F. Lagarrigue, A.R. Gingras, Z. Fan, K. Ley, and M.H. Ginsberg. 2018. Transmission of integrin beta7 transmembrane domain topology enables gut lymphoid tissue development. The Journal of cell biology. 217:1453–1465.

Tadokoro, S., S.J. Shattil, K. Eto, V. Tai, R.C. Liddington, J.M. de Pereda, M.H. Ginsberg, and D.A. Calderwood. 2003. Talin binding to integrin beta tails: a final common step in integrin activation. Science. 302:103–106.

Vranken, W.F., W. Boucher, T.J. Stevens, R.H. Fogh, A. Pajon, M. Llinas, E.L. Ulrich, J.L. Markley, J. Ionides, and E.D. Laue. 2005. The CCPN data model for NMR spectroscopy: development of a software pipeline. Proteins. 59:687–696.

Wegener, K.L., A.W. Partridge, J. Han, A.R. Pickford, R.C. Liddington, M.H. Ginsberg, and I.D. Campbell. 2007. Structural basis of integrin activation by talin. Cell. 128:171–182.

Ye, F., G. Hu, D. Taylor, B. Ratnikov, A.A. Bobkov, M.A. McLean, S.G. Sligar, K.A. Taylor, and M.H. Ginsberg. 2010. Recreation of the terminal events in physiological integrin activation. The Journal of cell biology. 188:157–173.

Zeiler, M., M. Moser, and M. Mann. 2014. Copy number analysis of the murine platelet proteome spanning the complete abundance range. Molecular & cellular proteomics : MCP. 13:3435–3445.

Zhu, L., J. Yang, T. Bromberger, A. Holly, F. Lu, H. Liu, K. Sun, S. Klapproth, J. Hirbawi,T.V. Byzova, E.F. Plow, M. Moser, and J. Qin. 2017. Structure of Rap1b bound to talin reveals a pathway for triggering integrin activation. Nature communications. 8:1744.

